# Non-invasive imaging of tau-targeted probe uptake by whole brain multi-spectral optoacoustic tomography

**DOI:** 10.1101/2021.07.10.451626

**Authors:** Patrick Vagenknecht, Maiko Ono, Artur Luzgin, Bin Ji, Makoto Higuchi, Daniela Noain, Cinzia Maschio, Jens Sobek, Zhenyue Chen, Uwe Konietzko, Juan Gerez, Riek Roland, Roger M. Nitsch, Daniel Razansky, Jan Klohs, Xose Luis Dean-Ben, Ruiqing Ni

**Author notes:** Corresponding author: Dr. Ruiqing Ni; Dr. Xose Luis Dean Ben, Address: Wagistrasse 12, 8952 Zurich, Switzerland.

## Abstract

**Aim:** Abnormal tau accumulation plays an important role in tauopathy diseases such as Alzheimer’s disease and Frontotemporal dementia. There is a need for high-resolution imaging of tau deposits at the whole brain scale in animal models. Here, we demonstrate non-invasive whole brain imaging of tau-targeted PBB5 probe in P301L model of 4-repeat tau at 130 μm resolution using volumetric multi-spectral optoacoustic tomography (vMSOT).

**Methods:** The binding properties of PBB5 to 4-repeat K18 tau and Aβ_42_ fibrils were assessed by using Thioflavin T assay and surface plasmon resonance assay. We identified the probe PBB5 suitable for vMSOT tau imaging. The imaging performance was first evaluated using postmortem human brain tissues from patients with Alzheimer’s disease, corticobasal degeneration and progressive supranuclear palsy. Concurrent vMSOT and epi-fluorescence imaging of *in vivo* PBB5 targeting (*i*.*v*.) was performed in P301L and wild-type mice. *Ex vivo* measurements on excised brains along with multiphoton microscopy and immunofluorescence staining of tissue sections were performed for validation. The spectrally-unmixed vMSOT data was registered with MRI atlas for volume-of-interest analysis.

**Results:** PBB5 showed specific binding to recombinant K18 tau fibrils, Alzheimer’s disease brain tissue homogenate by competitive binding against [^11^C]PBB3 and to tau deposits (AT-8 positive) in post-mortem corticobasal degeneration and progressive supranuclear palsy brain. *i*.*v*. administration of PBB5 in P301L mice led to retention of the probe in tau-laden cortex and hippocampus in contrast to wild-type animals, as also confirmed by *ex vivo* vMSOT, epi-fluorescence and multiphoton microscopy results.

**Conclusion:** vMSOT with PBB5 facilitates novel 3D whole brain imaging of tau in P301L animal model with high-resolution for future mechanistic studies and monitoring of putative treatments targeting tau.

## Introduction

The abnormal cerebral deposition of pathological tau fibrils is a characteristic feature of tauopathy-related neurodegenerative diseases including Alzheimer’s disease (AD), corticobasal degeneration (CBD), progressive supranuclear palsy (PSP) and parkinsonism linked to chromosome 17 ^1^. The microtubule-associated protein tau (MAPT) is located intracellularly and is composed of six isoforms classified into 4-repeat (4R) and 3-repeat (3R) species ^2^. Several tau positron emission tomography (PET) tracers have been developed, including the first generation [^18^F]flortaucipir ^3^, [^11^C]PBB3 ^4^, and [^11^C]THK5351, [^18^F]THK5117 ^5-7^; second generation [^18^F]MK-6240 ^8^, [^18^F]PM-PBB3 (APN1607) ^9^, [^18^F]JNJ-64326067 ^10^, [^18^F]RO948 ^11^, [^18^F]PI-2620 ^12^, and [^18^F]GTP1 ^13^. PET showed the spreading of tau in patients with AD, which correlates with axonal damage, neurodegeneration, functional network alterations, and cognitive impairment. Thereby, the tau bio-distribution represents a powerful bio-marker with great potential in disease staging ^14-23^. In addition, the tau tracer [^18^F]PM-PBB3 has been shown to facilitate detecting different pattern in patients with PSP and CBD compared to AD, indicating its capability for differential diagnosis ^9^.

Transgenic mouse models (mutations in the *MAPT* gene) recapitulate pathological features of tauopathy and have greatly advanced our understanding of disease mechanisms ^24-28^. *Ex vivo* high-resolution light-sheet microscopy with anti-tau antibodies or luminescent conjugated oligothiophenes (LCOs) enabled whole-brain mapping of tau bio-distribution and spread ^29-31^. However, capturing early tau deposits *in vivo* is needed for a better understanding of the link with other pathological alterations in the deep brain regions. *In vivo* microPET imaging of the cerebral tau accumulation in the transgenic tauopathy mouse has been achieved using [^18^F]PM-PBB3, [^11^C]PBB3, [^11^C]mPBB5 ^9, 32-35^, [^18^F]THK5117 ^36, 37^, [^18^F]JNJ-64349311 ^38^, and 4R-tau specific tracers [^18^F]CBD-2115 ^39^ and [^11^C]LM229 ^40^. PET is known to provide excellent accuracy to map the bio-distribution of tau in human subjects. However, microPET has a limited spatial resolution (0.7-1.5 mm) relative to the small mouse brain (∼10 mm^3^), which hinders accurate detection of tau, especially in small subcortical brain regions ^41^. Fluorescence tau imaging using PBB5 ^33, 42^, luminescent oligothiophene conjugated probe (h-FTAA) ^43^, BF-158 ^42^, Q-tau 4 ^44^, pTP-TFE ^45^, BODIPY derivative ^46, 47^ and fluorescent-labelled antibodies ^48^ has also been reported. However, fluorescence imaging only provides a planar view, limited depth and limited quantification capabilities. On the other hand, two-photon imaging of mice with a cranial window using HS-84 ^49^, methoxy-X04 ^50^, fluorescent-labelled antibodies ^51^ can follow the development of tau at cellular resolution, but with sub-millimeter field-of-view (FOV) and low penetration depth. Overall, existing imaging approaches are either limited by penetration depth or spatial resolution, which demands for non-invasive imaging tools providing high-resolution performance at whole-brain scales.

Recently, volumetric multi-spectral optoacoustic tomography (vMSOT) imaging has been shown to provide previously unavailable capabilities to visualize the bio-distribution of amyloid-β (Aβ) deposits in mouse models of AD amyloidosis (arcAβ and APP/PS1) ^52-54^. vMSOT capitalizes on the high sensitivity of optical contrast and the high resolution provided by ultrasound ^55, 56^, and can attain penetration depth of the whole mouse brain. State-of-the-art vMSOT embodiments enable whole-brain non-invasive imaging with 130 µm spatial resolution ^52, 53, 57-62^, i.e., almost an order of magnitude finer resolution compared to modern small-animal microPET scanners. In this study, we investigate on the capabilities of vMSOT assisted with the arylquinoline derivative PBB5 probe to enable *in vivo* high-resolution 3D transcranial mapping of tau across the entire mouse brain in 4R-tau P301L mouse models ^26^. The targeting performance of the PBB5 probe is further evaluated using *post-mortem* human brain tissues from patients with AD, PSP and CBD.

## Results

### Thioflavin T (ThT) and surface plasmon resonance (SPR) *in vitro* binding assay in recombinant fibrils

To characterize the absorbance spectrum, affinity, binding kinetics, and specificity of PBB5, ThT assay and SPR assay were performed using recombinant Aβ_42_, 4-repeat K18 tau fibrils. The Aβ_42_, 4-repeat K18 tau fibrils were validated using ThT and western blot (**Suppl. Fig. 1**). The results from *in vitro* ThT assay indicated that PBB5 binds to K18 tau with specific peak spectrum, although also binding to Aβ_42_ fibrils (**Fig. 1a-b**). We further characterized the kinetics and affinities of PBB5 to different fibrils using surface plasmon resonance binding assay.

**Fig. 1.**
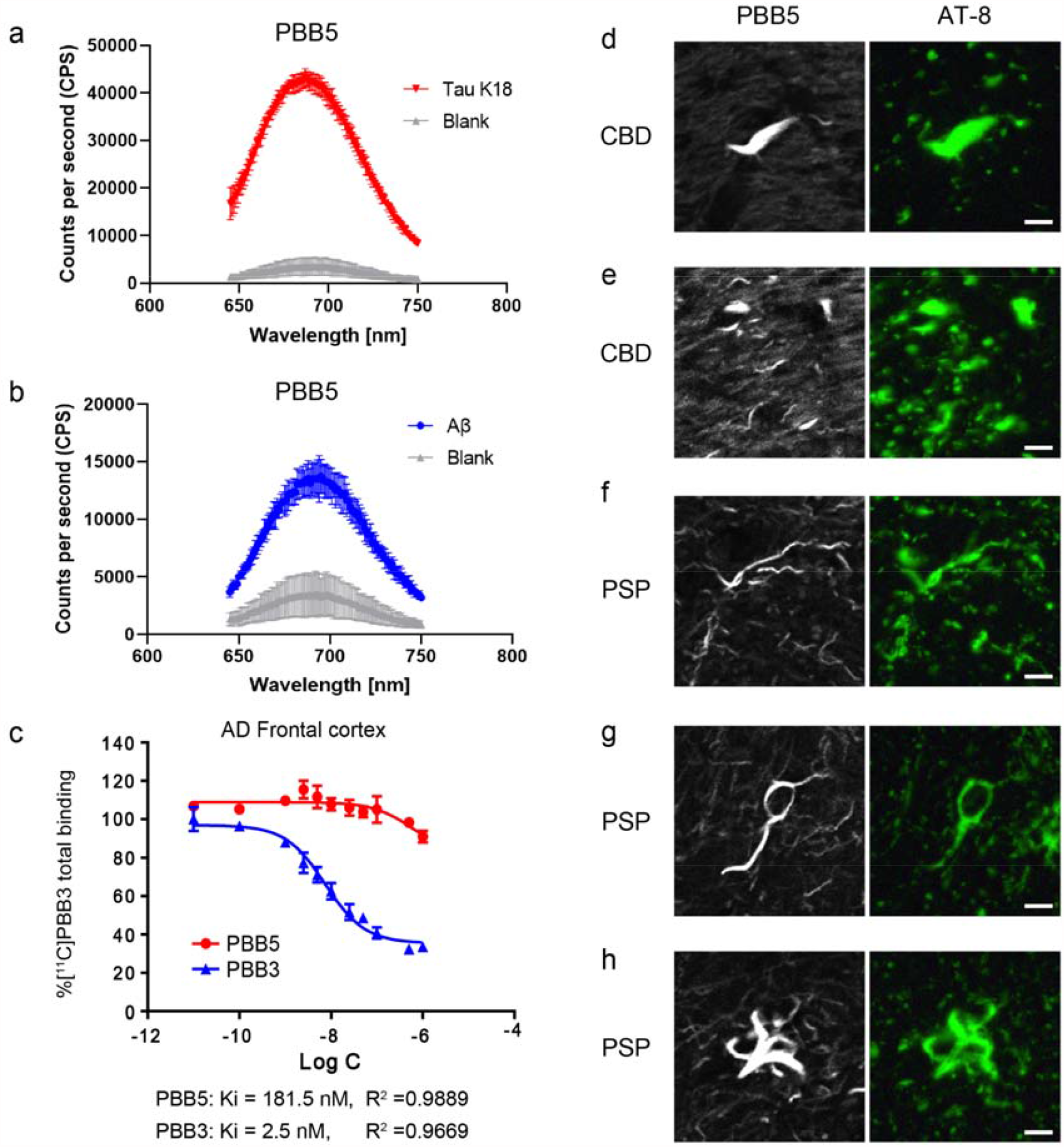
PBB5 characterization on recombinant fibrils and staining on human brain. (**a-b**) Thioflavin T binding assay using PBB5 on Aβ42, K18 4-repeat tau fibrils and blank (dd. water) with PBB5; (**c**) Binding of [^11^C]PBB3 in cortical homogenates derived from Alzheimer’s disease (AD). Total (specific + non-specific) binding of 5 nM of [^11^C]PBB3 in an AD temporal cortex sample blocked homologously by non-labelled PBB3 (blue) and heterologously by non-labelled PBB-5 (red). Inhibition of radioligand binding was described by 1-site model, and parameters resulting from curve fits are indicated; (**d-h**) PBB5-positive and AT-8-positive inclusions indicated coiled bodies (**d, g**), argyrophilic threads (**e, f**) accumulated in oligodendrocytes and tufted astrocyte (**h**) in the caudate/putamen from patients with corticobasal degeneration (CBD) and motor cortex from progressive supranuclear palsy (PSP); scale bar = 10 μm. AT-8: an anti-phosphorylated tau antibody.

### Binding assays and staining in human brain

We further characterized the binding property of PBB5 using brain tissues from patients with different tauopathies including AD brain tissue with mixed 3R, 4R-tau, CBD and PSP brain tissue with 4R-tau. Competitive binding assay in AD brain homogenates using different concentrations of unlabeled PBB5 and PBB3 against [^11^C]PBB3 (concentration: 5 nM, specific activity: 86.9 GBq/μmol, radiochemical purity: 96.7 %) indicated an inhibition constant (Ki) = 181.5 nM, and partial replacement for PBB5 (R^2^ = 0.9889, n = 4), compared to Ki = 2.5 nM for PBB3 (R^2^ = 0.9669, n = 4) (**Fig. 1d**). Staining using PBB5 and anti-phosphorylated tau antibody (AT-8) in the caudate/putamen from patients with CBD and motor cortex from PSP showed an overlapping signal, which indicates that PBB5 is capable of recognizing AT8 positive coiled body (**Figs. 1d, g**) and argyrophilic threads in oligodendrocytes (**Figs. 1e, f**), and tufted astrocyte (**Fig. 1h**).

### Non-invasive *in vivo* vMSOT of PBB5 uptake in the mouse brain

The absorption spectrum of PBB5 expands within the far-red range (∼590-690 nm, **Fig. 1a**), where light penetration is significantly enhanced with respect to that achieved for shorter wavelengths. This facilitates distinguishing the bio-distribution of PBB5 from endogenous chromophores deoxyhemoglobin (Hb), and oxyhemoglobin (HbO) via spectral unmixing of vMSOT images acquired *in vivo*. The surface-weighted PBB5 bio-distribution was also measured in the epi-fluorescence mode in both P301L and wild-type mice by means of concurrent planar fluorescence-vMSOT system. The set-up includes a custom-build hybrid imaging system with a fiberscopic insert (**Fig. 2b**) as described in detail elsewhere ^52, 53^. The vMSOT imaging data analysis pipeline consisted on the following steps. First, 3D optoacoustic images were reconstructed for multiple excitation wavelengths (**Fig. 2c**). Then, spectral unmixing was performed to isolate the bio-distributions of HbO and PBB5. Finally, co-registration with a MRI mouse brain atlas ^63^ for volume-of-interest (VOI) analysis was performed (**Fig. 2d**). After *i*.*v*. bolus injection of PBB5 in mice through the mouse tail vein (n = 20 in total), an increase in the fluorescence and/or spectrally unmixed PBB5 signal was observed in the mouse brain parenchyma, arguably indicating that the probe passed the blood-brain barrier (BBB). Epifluorescence images of the brain corroborated the increase in signal associated to PBB5, albeit providing no depth information and significantly inferior resolution compared to vMSOT (**Fig. 2e**). A 3D view of the unmixed image for PBB5 distribution corresponding to 60 minute post-injection is shown in **Suppl. Fig. 1** and **Suppl. Video 1, 2**.

**Fig. 2.**
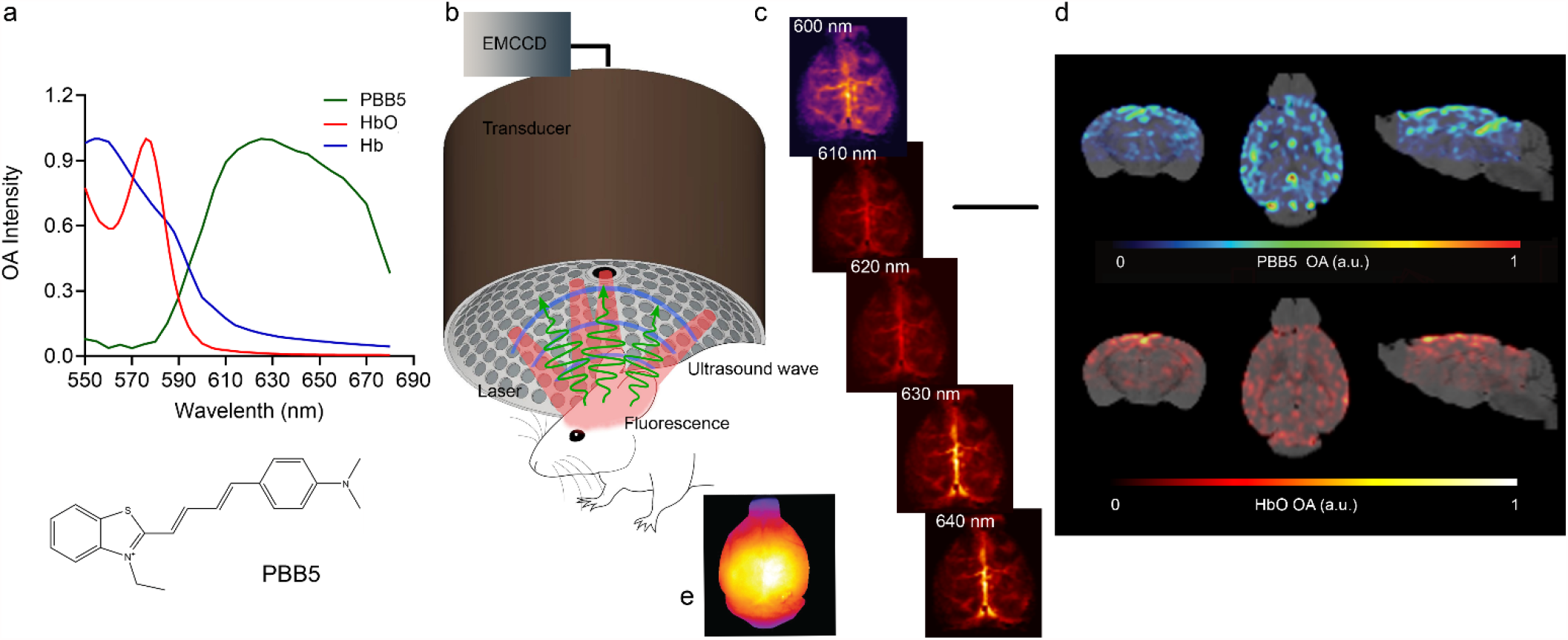
Non-invasive tau vMSOT imaging pipeline. (**a**) Chemical structure of the probe PBB5 and extinction spectrum of HbO and Hb along with the spectrum of PBB5 measured by volumetric multi-spectral optoacoustic tomography (vMSOT); (**b**) Set-up of the vMSOT system for tau mapping across entire mouse brain; (**c**) Volumetric reconstructions of the *in vivo* vMSOT data for five distinct excitation wavelengths (600, 610, 620, 630, 640 nm) used for spectral unmixing; (**d**) coronal, horizontal and sagittal view of PBB5 and HbO; Absorbance intensity scale: 0-1. (**e**) Simultaneous epi-fluorescence imaging in one P301L moues brain after *i*.*v*. injection of PBB5.

### Spectral unmixing of the vMSOT data

Spectral unmixing can generally isolate the bio-distribution of any spectrally-distinctive probe from endogenous absorbers in biological tissues. However, spectral coloring effects associated to wavelength-dependent attenuation of light lead to cross-talk artefacts when considering the theoretical spectra of the absorbing substances present in the sample ^64, 65^. This is particularly important for spectral windows exhibiting sharp variations of the hemoglobin absorption, e.g. around the 600-630 nm wavelengths (**Fig. 2a**) ^66^. Herein, we consider the sequence of images taken during injection of PBB5 for a mouse in which no motion was detected to optimize the multi-spectral unmixing procedure. Specifically, the wavelengths and absorbing components were optimized so that the unmixed bio-distribution of PBB5 matches that obtained by subtracting a reference image taken before injection for the sequence vMSOT images taken at 640 nm wavelength. Note that this approach is only applicable when such a reference is available, which is generally not the case due to sample motion e.g. during i.v. injection. We found that the unmixing performance was optimal when considering five wavelengths (600, 610, 620, 630 and 640 nm) and only oxygenated hemoglobin (HbO) and PBB5 as absorbing components. **Figs. 3a-c** show the unmixed bio-distributions of HbO and PBB5 in this case for a time point ∼100 seconds after injection. The unmixed bio-distribution of PBB5 is shown to match the differential (baseline-subtracted) vMSOT image at 640 nm (**Fig. 3d**). The good matching between the two approaches is better observed by considering the time profiles of selected time points (superior sagittal sinus, superficial vessels and deep brain region) in the unmixed and differential vMSOT images (**Fig. 3)**. This corroborates the validity of multispectral unmixing with the selected wavelengths and components as a method to isolate the bio-distribution of PBB5.

**Fig. 3.**
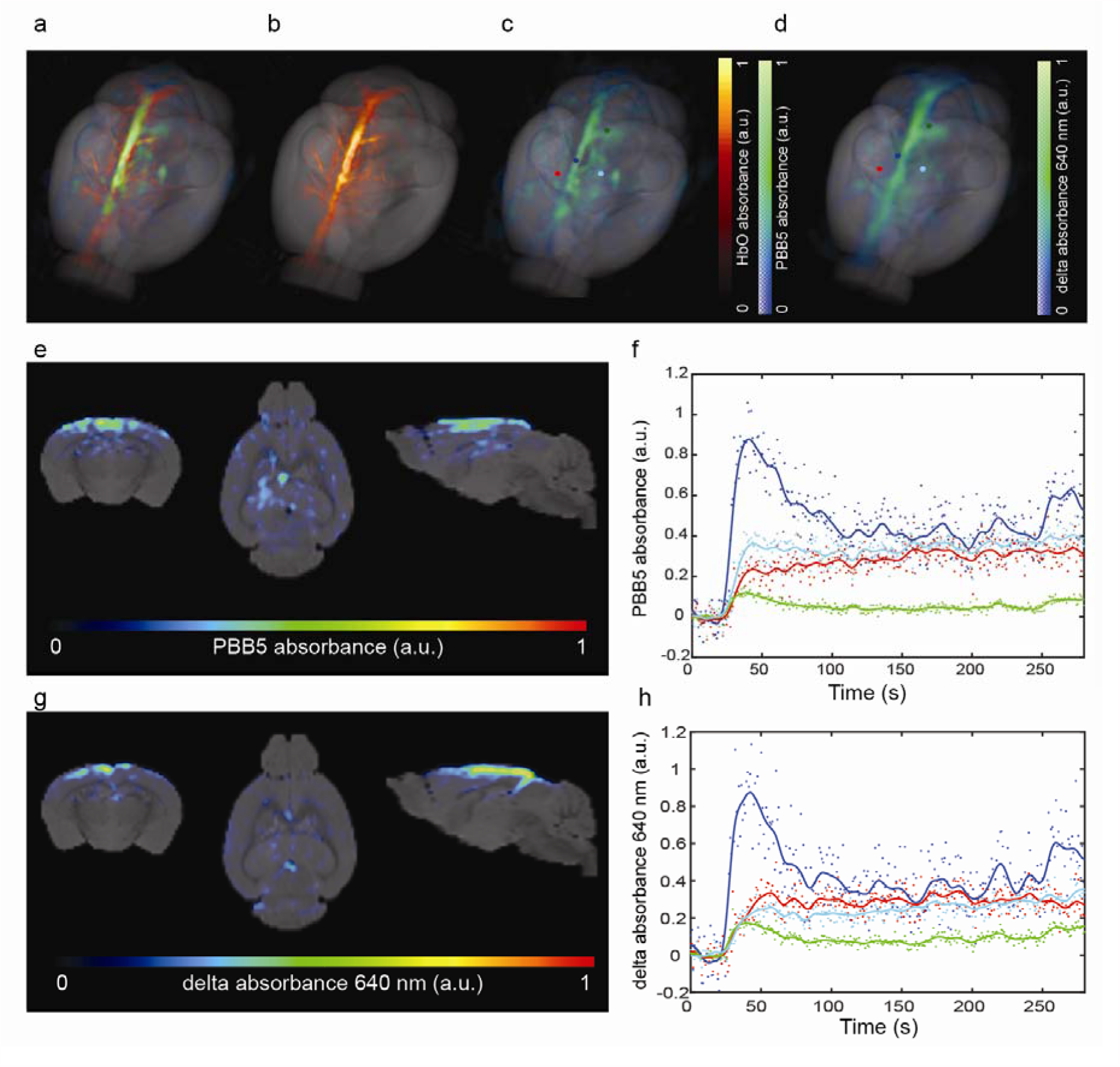
Comparison of different vMSOT processing methods. (**a-d**) 3D rendering of vMSOT data in mouse brain unmixed for PBB5 distribution (c), Image acquired at 600 nm excitation wavelength reveals the major cerebral vessels (b), and overlay (a); (d) Baseline-subtracted single wavelength vMSOT image acquired at 640 nm; (**e, g**) vMSOT unmixed PBB5 absorbance image and baseline-subtracted single wavelength vMSOT image acquired at 640 nm. PBB5 was injected i.v. at 30 s. Coronal, sagittal and horizontal views overlaid over the masked magnetic resonance imaging-based brain atlas. (**f, h**) Time-lapse curves of unmixed PBB5 absorbance vMSOT image (f), and baseline-subtracted single wavelength vMSOT image (h) acquired at 640 nm in the cortical and deep brain regions; The temporal evolution of the unmixed AOI987 absorbance and baseline-subtracted signals was analyzed in four different brain regions: cortex (red), superior sagittal sinus (dark blue), hippocampus (green), vessel (light blue) on the cortical surface indicated in (**c, d**)

### Dosage-dependent performance

The optimal dosage of PBB5 to be injected i.v *in vivo* so that it can be clearly detected in the vMSOT images was established by testing different concentrations of PBB5 (5, 25, 50 mg/kg weight) in P301L and wild-type mice. **Figure 4** showed the representative pattern using different dosage of PBB5 in P301L mouse brain. The mice were scanned *in vivo* with vMSOT at pre-injection, during injection (7 minute dynamic scan), and at 20, 40, 60, 90 and 120 minute following *i*.*v*. injection of PBB5. A dependence on the unmixed PBB5 signal in the vMSOT images with the concentration of the probe was clearly observed at 20-60 min post-injection (**Fig. 4**). The time profiles at approximate locations in the superior sagittal sinus further indicate a clear signal enhancement with concentration. Due to the abundant endogenous signal Hb/HbO in the mouse brain, negligible signal increase was detected using for 5 mg/kg dose. *i*.*v*. injection of 25 mg/kg PBB5 provided sufficient vMSOT signal increase to allow detecting differences in the unmixed images (**Fig. 4b**).

**Fig. 4.**
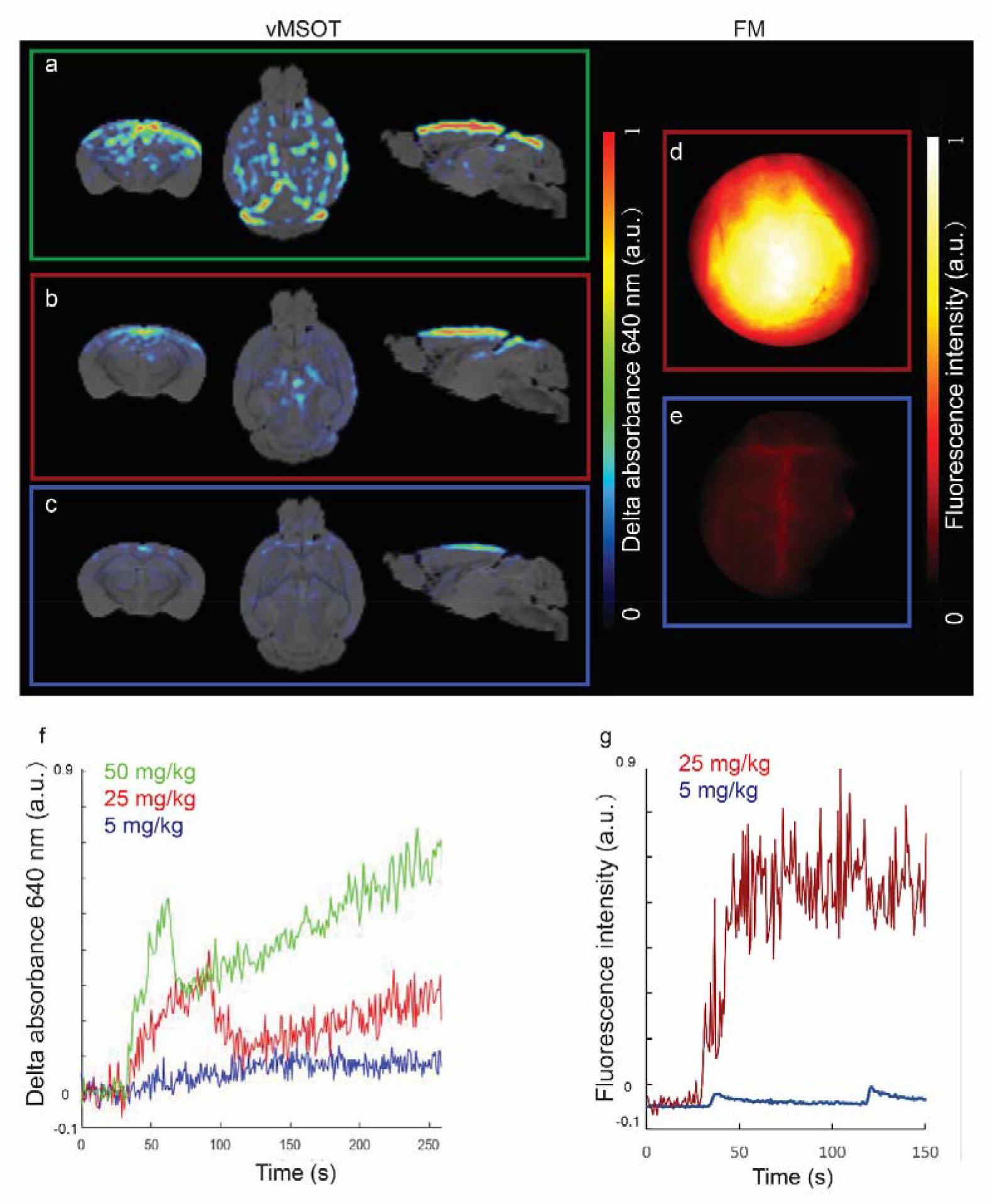
Dose determination for *in vivo* tau imaging with vMSOT. (**a-e**) vMSOT images of three different concentration of PBB5, 5 mg/kg weight (blue square, c), 25 mg/kg weight (red square, b), and 50 mg/kg weight (green square, a) and epifluorescence images of 5 mg/kg weight (blue square, e), 25 mg/kg weight (red square, d); (**f)** Time curve of unmixed PBB5 absorbance profile during the first 300 seconds (within 7 minute dynamic *i*.*v*. injection using three different concentration of PBB5, 5 mg/kg weight (blue line), 25 mg/kg weight (red line), 50 mg/kg weight (green line); No clear signal increase was detected using 5 mg/kg weight dose; (**g**) Fluorescence intensity curve of PBB5 using 5 mg/kg weight (light blue) and 25 mg/kg weight (dark blue). PBB5 was injected *i*.*v*. at 30 s.

Results from fluorescence imaging indicated a similar dose-dependent signal (**Fig. 4**). The fluorescence signal is very intense already at 25 mg/kg of PBB5, and sufficient fluorescence signal increase can be detected using *i*.*v*. injection of 5 mg/kg PBB5 (not detectable by vMSOT). Similar to the pattern in the spectrally unmixed vMSOT signal intensity. As the intensity of PBB5 absorbance and fluorescence intensity was stable from 40-60 minute and slight decrease upto 120 minute, we chose 60 minute scanning time frame.

### PBB5 bio-distribution in P301L and wild-type mice

P301L (n = 3) and wild-type mice (n = 3) were imaged at different time points before, during and after injection of PBB5 (25 mg/kg weight *i*.*v*.) using the vMSOT system. The unmixed images for the PBB5 channel were superimposed onto the MRI atlas for VOI analysis (**Fig. 5a**). The time courses of PBB5 absorbance in different brain regions of P301L and wild-type mice were assessed. In addition we used cerebellum which is void of tau accumulation for P301L model as reference brain region, Two-way ANOVA with Bonferroni *post hoc* analysis showed the interaction between brain region and genotype. A significantly higher PBB5 retention (average of absorbance at 60 minute post-injection) was observed in the cortex, hippocampus and thalamus of P301L mice compared to wild-type mice (**Fig. 5**). Consistently, the PBB5 absorbance signals were higher in the cortex, hippocampus and thalamus of P301L mice compared to their respective wild-type mice. Similar temporal profiles of vMSOT and planar fluorescence signals were observed throughout the cortical region **(Figs. 5**). Robust correlation was observed between fluorescence and unmixed vMSOT PBB5 absorbance signal (p < 0.0001, Pearson’s rank correlation analysis (**Fig. 4**). Next we calculated using cerebellum as a reference brain region, Similar patten was observed. Consistently, the PBB5 absorbance ratio were higher in the cortex/cerebellum, hippocampus/cerebellum and thalamus/cerebellum of P301L mice compared to their respective wild-type mice. The test-retest correlation analysis between independent analysis was shown in **Suppl. Fig. 3** indicating the repeatability of the VOI analysis (interrater variability and intrarater variability).

**Fig. 5.**
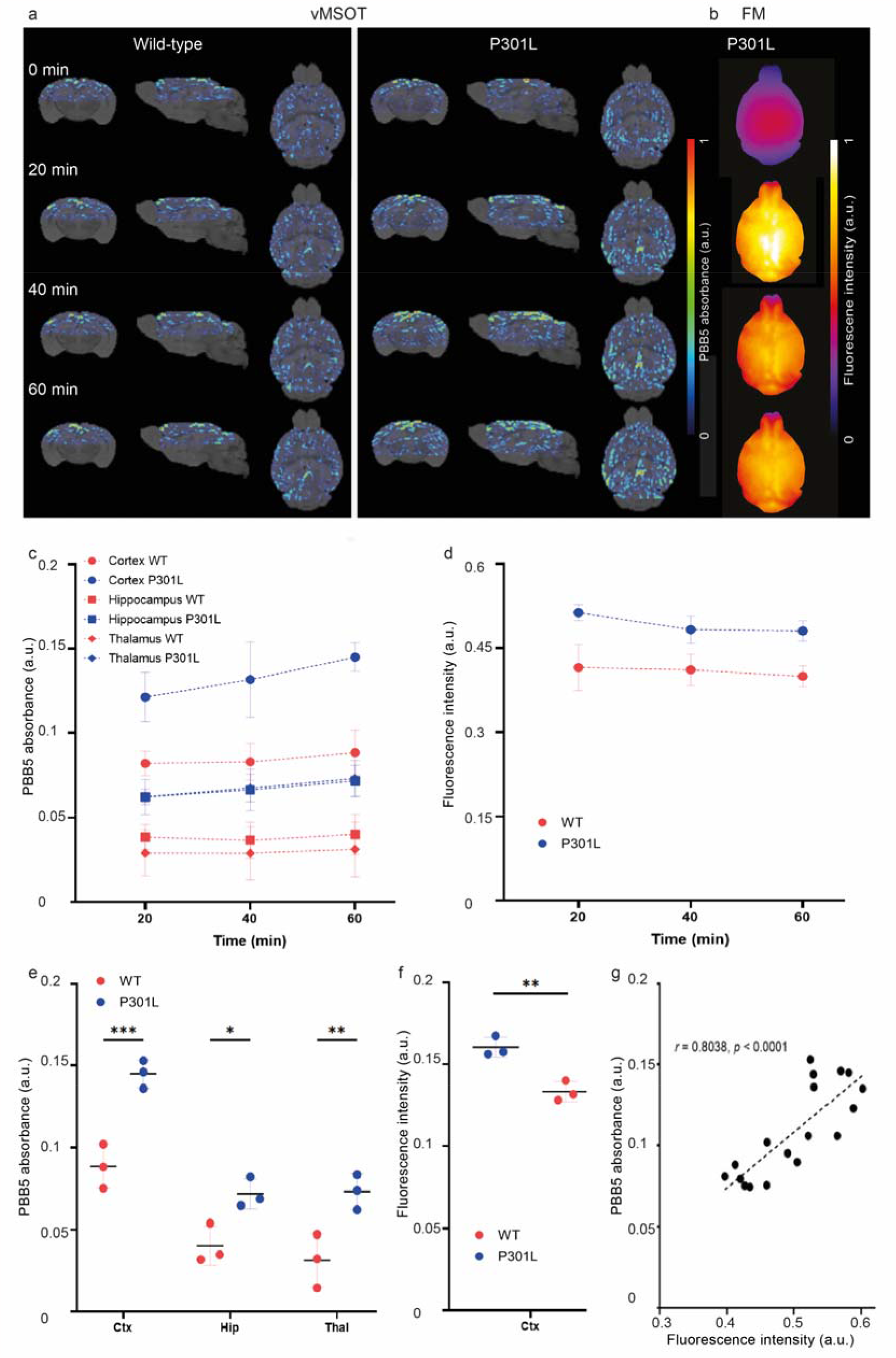
Regional tau distribution revealed by *in vivo* vMSOT imaging using PBB5 probe in P301L and wild-type mice, (**a**) Wild-type (WT) and transgenic P301L mice; at pre-injection, 20, 40, 60 min following dye administration showing coronal, sagittal and horizontal views overlaid over the masked magnetic resonance imaging-based brain atlas. PBB5 absorbance signal strength is indicated by rainbow color-map; (**b**) Example of epi-fluorescence images from one one P301L mouse at 20, 40, 60 min following dye administration; (**c, d**) Time course of cortical, hippocampal, thalamic volume-of-interest PBB5 signal (absorbance signal) and cortical region-of-interest fluorescence intensity; (**e, f**) regional comparison of probe absorbance signal retention and fluorescence intensity at 60 min post-injection, Data are presented as mean±SD; P301L (n = 3), and NTL (n = 3); *p<0.05, **p<0.01, ***p<0.001 comparison between WT and P301L mice. Cortex: Ctx; Hippocampus: Hip; Thalamus: TH; (**g**) Correlation between optoacoustic and Fluorescence imaging across different mice using Pearson rank analysis.

### *Ex vivo* validation

To validate the *in vivo* imaging results, the mouse brains were dissected after *in vivo* imaging and imaged *ex vivo* using the same vMSOT set-up. **Figs. 6** displays a comparison of the 3D unmixed PBB5 signal overlaid with the MRI from a representative P301L mouse. The accumulation of PBB5 signal in the cortex and the hippocampus of P301L mouse suggest specific binding of the probe to these regions known to express high tau load. Imaging on coronal brain slices (∼2 mm thickness, coronal slices cut using a brain matrix at Bregma 0- -2 mm) indicate retention of signal in the brain of P301L mouse.

**Fig. 6.**
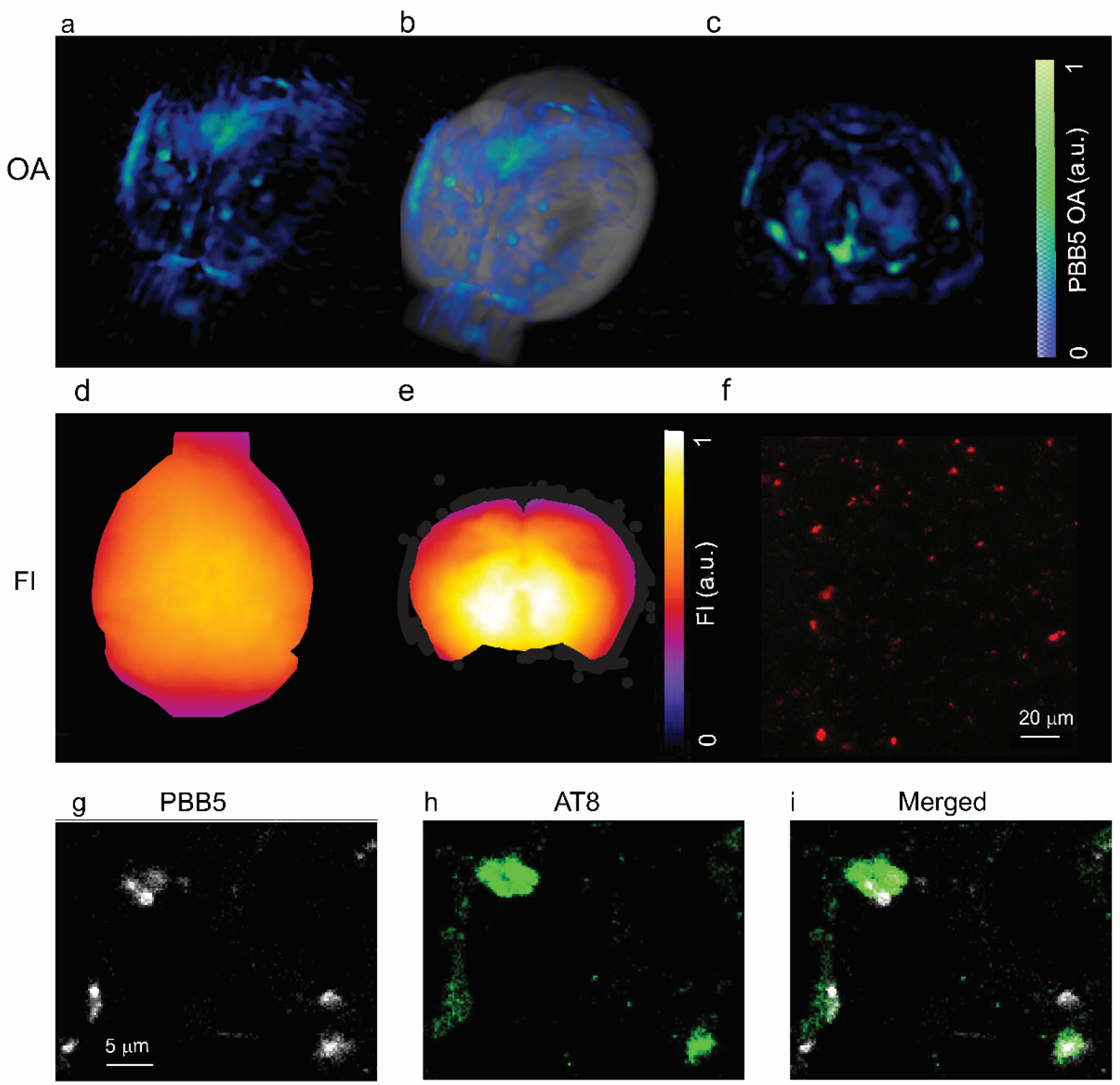
*Ex vivo* validation. **(a-c)** *Ex vivo* vMSOT of whole brain, and brain slice at 90 minute after PBB5 i.v. injection; (**a**) 3D rendering of ex vivo vMSOT data unmixed for PBB5 distribution in P301L mouse brain; **(b**) Overlay of (a) on MRI structural data; (**c**) *ex vivo* vMSOT of 1 mm mouse brain slice data unmixed for PBB5 distribution in P301L mouse brain. PBB5 absorbance signal strength is indicated by blue-green color-map; (**d, e**) Epi-fluorescence of (**a, c**); (**f**) Multiphoton microscopy (MPM) regional quantification multiphoton. Scale bar = 20 μm; (**g-i**) Confocal microscopic images of hippocampus sections from P30L mice. PBB5 (white), Alexa488-AT-8 (green) in the hippocampus areas. Scale bar = 5 μm;

To validate the *in vivo* PBB5 signal distribution in deep brain regions imaged with vMSOT, we imaged fixed brain from P301L and wild-type mice (transcardiac perfusion and 4 % PFA fixation) using Leica TCS SP8 multiphoton microscopy at 20× magnification. Lambda scans was performed on the suspected tau deposits, and showed fluorescence emission spectrum indicating PBB5 (**Fig. 6**). This result further demonstrated that PBB5 efficiently binds to tau *in vivo*. In congruence with the *in vivo* imaging findings, tau deposits morphology was clearly observed in tissue slices with stronger PBB5 signal found in the cortex of P301L mice (**Fig. 6**). Immunofluorescence staining performed on horizontal brain tissue sections from P301L and wild-type mice co-staining with AT-8 antibody (**Fig. 6, Suppl. table 1**).

## Discussion

New tools for non-invasive mapping of tau deposits with high-resolution in animal models of tauopathy are imperative for understanding the spreading of tau deposits ^67^ and for translational development of tau-targeted therapeutic and diagnostic tools ^68, 69^. Herein, we identified PBB5 as a suitable tau imaging probe for vMSOT that binds with high sensitivity and specificity to tau aggregates. This was used to establish a novel *in vivo* transcranial vMSOT imaging approach to map whole brain tau deposits at 130 μm resolution in a P301L mouse model.

The criteria for selecting an appropriate tau-specific probe for vMSOT imaging include suitable absorption spectrum to allow unambiguous unmixing from the endogenous signal of blood (preferably with peak absorption at > 600 nm optical wavelength), high-affinity, low toxicity, low non-specific binding, photostability, low toxicity as well as low molecular weight and suitable lipophilicity to allow sufficient blood-brain barrier passage, and biocompatibility ^70^. Herein, we chose PBB5 with a preference to 4R-tau >10 times higher than to Aβ_42_ fibrils. Its peak absorption at 630-640 nm, where the absorption of hemoglobin decays, further facilitates distinguishing it from blood. Our ThT assay and SPR assay showed a specific binding of PBB5 to recombinant K18 tau, higher than to binding to Aβ_42_ fibrils. The binding affinity and kinetics (Kon, Koff) of PBB5 is suitable for *in vivo* imaging.

A competitive binding assay against [^11^C]PBB3, PBB5 was further shown to have an affinity Ki of 181 nM in *post-mortem* brain tissue from patients with AD cortex. The binding affinity is in line with the previously reported affinity of PBB5 ^33^. Although the specificity and brain penetration of PBB5 is lower than that of PBB3 (with peak absorption at 405 nm) ^33^ or PM-PBB3 (emission at 525 nm) ^9^, its near-infrared absorption spectrum allows for epi-fluorescence and vMSOT imaging of deep brain regions. Staining with PBB5 and AT-8 of brain tissues from caudate/putamen patients with CBD and motor cortex from PSP showed an overlapping signal demonstrating that PBB5 is capable of recognizing tau accumulation in coiled body and argyrophilic threads inside oligodendrocytes in brain from CBD and PSP, as well as tufted astrocytes in brain from PSP.

Tau plays an important role in the pathogenesis of AD and other primary tauopathy diseases such as CBD and PSP ^29, 71, 72^. Ongoing clinical trials targeting at reducing tau have shown promising results. These include antibodies gosuranemab BIIB092 or non-pharmacological treatments ^73-77^. Tau imaging has however been challenging due to the structural diversity of tau isoforms, the difference between 4R and 3R tau, its intracellular location, as well as the specificity and off-target binding of tau imaging probes ^78, 79^. PET assisted with the tau tracer [^18^F]PM-PBB has been shown to detect different patterns in patients with PSP and CBD compared to AD, indicating a role in differential diagnosis ^9^. Recent cryo-EM has shown that PM-PBB3 binds to tau fibrils in AD brain ^80^. An *In silico* study reported THK5351 probes, T807 binding to different sites on tau fibrils ^4, 81^ as well as off-target binding sites ^79^. Previous autoradiography and PET studies indicated that PBB analogs, THK5351 or THK5117 and JNJ-64349311 but not T807 can detect tauopathy in tau mouse models (P301L, PS19 line) ^32-34, 36, 38, 82, 83^.

In P301L (CaMKII) mice, tau deposits start at 5 months-of-age, first in the limbic system (entorhinal cortex and hippocampus) and subsequently spreading to the neocortex ^26, 84^. Tauopathy deposits in P301L (Thy1.2) mice ^26^ are most pronounced in the cortex, amygdala and hippocampus, moderate in the brain stem and striatum, and negligible in the cerebellum. Thus we chose cerebellum as reference brain region. Similar to PBB3 and PM-PBB3, PBB5 detects the AT-8 stained neurofibrillary tangle, ghost tangles, tau deposits in astrocytes and oligodendrocytes in the brain from PSP, CBD ^85^. In P301L as well as in other tauopathy mouse models, the neurofibrillary tangle is rear and less fibrillar structure is present in the mouse brain ^26, 34, 84^. The cortical and hippocampal signals detected by vMSOT *in vivo* and *ex vivo* using PBB5 are in accordance with immunofluorescence staining results, and with the known tau distribution in the P301L mouse brain ^84, 86^. NIRF imaging using PBB5 and PET using [^11^C]mPBB5, respectively, have been previously reported for mapping tau deposition in the brain stem and spinal cord of P301S mice ^33^. However, NIRF imaging detection in deep brain regions was hindered by strong absorption and scattering of the excitation light and emitted fluorescence. Sub-millimeter scale intravital microscopy enables the visualization of tau deposits, but is highly invasive and can only cover a very limited FOV ^49^. We recently reported on large FOV fluorescence microscopy imaging of tau in P301L mice with 6 micron resolution, which however only provided a planar view ^43^. As the spatial resolution of vMSOT is not altered by photon scattering but rather governed by ultrasound diffraction, it enables high-resolution mapping and quantification of endogenous tissue chromophores or spectrally distinctive exogenous probes at millimeter to centimeter scale depths ^55, 87, 88^.

There are several limitations in the current study that need to be highlighted. We did not take into account the spectral colouring effect associated to wavelength-dependent optical attenuation, which may cause distortion in the vMSOT spectra rendered from deep locations ^65, 89^. These factors may lead to cross-talk artefacts in the unmixed images corresponding to the contrast agent. Advanced algorithms are required for attaining more accurate performance ^89^. In addition, future longitudinal studies are required to determine the sensitivity and specificity of the proposed methodology, how early PBB5 positive tau can be detected, and whether it can follow the spreading of tau in the brain ^90^.

In conclusion, we demonstrated non-invasive whole-brain high-resolution imaging of tau in P301L mice with a state-of-the-art vMSOT system, which is not feasible with other imaging modalities. This platform provides new tool to study tau spreading and clearance in tauopathy mouse model, foreseeable in monitoring of tau targeting therapeutics.

## Materials and methods

### Animal model

Mice transgenic for *MAPT P301L*, overexpressing the human 2N/4R tau under neuron-specific Thy1.2 promoter (pR5 line, C57B6.Dg background) ^26, 43, 86, 91, 92^, and wild-type littermate mice were used (18 months-old, P301L n = 10, wild-type n = 10, both genders). Of which, N = 7 from each group was used in initial testing and for different dosage (**Fig. 4**). N = 3 from each group was used for **Fig. 5**. Animals were housed in individually ventilated cages inside a temperature-controlled room, under a 12-hour dark/light cycle. Pelleted food (3437PXL15, CARGILL) and water were provided *ad-libitum*. All experiments were performed in accordance with the Swiss Federal Act on Animal Protection and were approved by the Cantonal Veterinary Office Zurich (permit number: ZH082/18, ZH162/20).

### Post-mortem human brain tissues

*Post-mortem* human brains were obtained from autopsies carried out at the Center for Neurodegenerative Disease Research of the University of Pennsylvania Perelman School of Medicine on patients with AD, CBD and PSP. Tissues for homogenate binding assays were frozen, and tissues for histochemical and immunohistochemical labeling were fixed in 10 % neutral buffered formalin followed by embedding in paraffin blocks. All procedures involving the use of human materials were performed in accordance with the ethical guidelines of the Institutional Review Boards (IRBs) of the University of Pennsylvania, and the National Institutes for Quantum and Radiological Science and Technology.

### *In vitro* [^11^C]PBB3 radiosynthesis and binding assay

Frozen tissues derived from the frontal cortex of an AD patient were homogenized in 50 mM Tris-HCl buffer, pH 7.4, containing protease inhibitor cocktail (cOmpleteTM, EDTA-free; Roche), and stored at -80 °C until analyses. [^11^C]PBB3 was synthesized as described previously^85^. To assay radioligand binding with homologous or heterologous blockade, these homogenates (100 µg tissue) were incubated with 5 nM [^11^C]PBB3 (specific radioactivity: 86.9 GBq/µmol) in the absence or presence of non-radiolabeled PBB3 or PBB5 at varying concentrations ranging from 1×10^−11^ to 5×10^−7^ M in Tris-HCl buffer containing 10 % ethanol, pH 7.4, for 30 minute at room temperature. Non-specific binding of [^11^C]PBB3 was determined in the presence of 5×10^−7^ M PBB3. Samples were run in quadruplicate. Inhibition constant (Ki) was determined by using non-linear regression to fit a concentration-binding plot to one-site and two-site binding models derived from the Cheng-Prusoff equation with GraphPad Prism version 5.0 (GraphPad Software), followed by F-test for model selection.

### Immunohistochemical staining on *post-mortem* brain tissues from patients with CBD and PSP

For fluorescence labeling with PBB5, deparaffinized sections were incubated in 50 % ethanol containing 2 µM of PBB5 at room temperature for 30 minutes. The samples were rinsed with 50% ethanol for 5 minutes, dipped into distilled water twice for 3 minutes, and mounted in non-fluorescent mounting media (VECTASHIELD; Vector Laboratories). Fluorescence images were captured using an FV-1000 confocal laser scanning microscope (Olympus, excitation at 635 nm and emission at 645-720 nm). Following fluorescence microscopy, all sections were autoclaved for antigen retrieval and immunohistochemical stained with anti-phosphorylated antibodies AT-8 (pSer202/pThr205, MN1020, Invitrogen, 1:250). Immunolabeling was then examined using a DM4000 microscope (Leica, Germany).

### *In vivo* imaging with the hybrid fluorescence and vMSOT system

Simultaneous vMSOT and planar fluorescence imaging at pre-, during, and post *i*.*v*. bolus injection of PBB5 was performed using a previously established hybrid fluorescence and vMSOT system ^52, 53, 93-95^, consisting of an epi-fluorescence fiberscope and a vMSOT system capable of covering the entire brain. The FOV provided is approximately 10×10 mm^2^ for epi-fluorescence imaging and 15×15×15 mm^3^ for vMSOT imaging, while the achievable spatial resolution is approximately 40 μm and 130 μm for epi-fluorescence and vMSOT, respectively ^53, 59, 88, 96^. Mice were first anesthetized with an initial dose of 4 % isoflurane (Abbott, Cham, Switzerland) in an oxygen/air mixture (200/800 mL/minute), and subsequently maintained at 1.5 % isoflurane in oxygen/air (100/400 mL/minute) throughout the measurement. The fur and the scalps over the head of the mice were then removed. The mice were placed in prone position on a heating pad with feedback control to maintain a constant body temperature. The mice were subsequently injected with a 100 μl bolus containing PBB5 (**Fig. 2**, dissolved in dimethyl sulfoxide (DMSO), 0.1 M PBS pH 7.4) through the tail vein. To establish the optimal dosage four P301L and four wild-type mice were used for dose response experiment (5, 25, 50 mg/kg weight). In the subsequent experiment the dose of 25 mg/kg body weight is chosen and used in the following experiment. For vMSOT, the pulse repetition frequency of the laser was set to 25 Hz and the laser wavelength tuned between 550 and 660 nm (5 nm step) on a per pulse basis. Epi-fluorescence imaging was performed by coupling the same beam from the pulsed OPO laser into the excitation fiber bundle. The excited fluorescence field was collected by an imaging fiber bundle comprised of 100,000 fibers and then projected onto an EMCCD camera (Andor iXon life 888, Oxford Instruments, UK). vMSOT and epi-fluorescence signals were recorded simultaneously before injection (108 s duration), during injection (432 s duration with *i*.*v*. injection starting at 30 s after the beginning of acquisition) and 20, 40, 60, 90 and 120 minute post-injection (108 s duration each).

### vMSOT image reconstruction and multi-spectral analysis

During the experiments, vMSOT images were reconstructed in real-time by using a graphics processing unit (GPU)-based implementation of a back-projection formula ^52, 53, 97^. The reconstructed images were further processed off-line to unmix the bio-distribution of PBB5 ^53^. Specifically, per-voxel least square fitting of the spectral signal profiles to a linear combination of the absorption spectra of oxygenated hemoglobin (HbO) and PBB5 was performed. Wavelengths between 600 and 640 nm (10 nm step) were considered. The optimum wavelengths and unmixing components were determined by comparing the unmixed bio-distribution of the probe with that obtained by the pre-injection image during injection of the probe. It was found that including deoxygenated hemoglobin led to larger errors in the bio-distribution of the probe.

The probe absorption spectra was experimentally determined as the average spectra of the differential (baseline-subtracted) vMSOT image during bolus perfusion at several major vessels in the brain. The vMSOT spectrum of PBB5 approximately matched the absorption spectrum measured with a spectrophotometer (Avantes BV, Apeldoorn, The Netherlands). The absorption spectrum of HbO was taken from an online database ^66^. The effective attenuation coefficient was estimated by considering a constant reduced scattering coefficient of 10 cm-1 for all mice and an optical absorption coefficient corresponding to the unmixed bio-distribution of blood and PBB5.

### Co-registration with MRI atlas and VOI analysis of the vMSOT data

Registration between vMSOT and MRI/atlas provides anatomical reference for regional analysis ^53, 98, 99^. These images were co-registered with T_2_-weighted structural MRI images (Ma-Benveniste-Mirrione-T_2_ ^63^) in PMOD 4.2 (Bruker, Germany) by two readers independently. VOI analysis of 15 brain regions was performed using the embedded Mouse VOI atlas (Ma-Benveniste-Mirrione) in PMOD ^53^. Specifically, dynamic time course and retention (60 min) of regional PBB5 absorbance intensity (a.u.) were calculated. Extra-cranial background signal was removed with a mask from the VOI atlas.

#### *Ex vivo* vMSOT imaging

To validate the *in*- and *ex vivo* signal, one P301L mice were perfused under ketamine/xylazine/acepromazine maleate anesthesia (75/10/2 mg/kg body weight, *i*.*p*. bolus injection) with ice-cold 0.1 M PBS (pH 7.4) and in 4 % paraformaldehyde in 0.1 M PBS (pH 7.4), and fixed for 4 h in 4 % paraformaldehyde (pH 7.4) and then stored in 0.1 M PBS (pH 7.4) at 4 °C. The dissected brain was imaged using vMSOT and hybrid epifluorescence imaging. The brain was cut coronally using a mouse brain matrix (World precision medicine, US) into 2 mm thickness at approximately 0 mm- -2 mm, and imaged again using the same set-up. For this, the spherical array was positioned pointing upwards and filled with agar gel to guarantee acoustic coupling, which served as a solid platform to place the excised brain and brain slice. Uniform illumination of the brain surface was achieved by inserting three arms of the fiber bundle in the lateral apertures of the array and a fourth one providing light delivery from the top. All recorded OA signals were normalized with the calibrated wavelength-dependent energy of the laser pulse. The bio-distribution of the probe was estimated via multi-spectral unmixing considering the vortex component algorithm (VCA) considering optical wavelengths from 600 to 655 nm (5 nm step) ^100, 101^.

### *Ex vivo* immunofluorescence and confocal imaging

Horizontal brain sections (40 μm) were cut and co-stained with PBB5 and anti-phosphorylated tau (pSer202/pThr205) antibody AT-8 (details in **Suppl. Table 1**). Sections were counterstained using DAPI ^86^. The brain sections were imaged at × 20 magnification using Axio Oberver Z1 and at × 63 magnification using a Leica SP8 confocal microscope (Leica, Germany) for co-localization of PBB5 with AT-8. The images were analyzed using ImageJ (NIH, U.S.A).

### *Ex vivo* multiphoton microscopy

Fixed brains from one P301L and one wild-type mice were imaged at ×20 magnification using Leica TCS SP8 Multiphoton microscopy and analyzed using ImageJ (NIH, United States). Lambda scan 3D rendering Identical settings resolution with Z stack and gain were used.

### Statistics

Group comparison of PBB5 absorbance in multiple brain regions at different time points was performed by using two-way ANOVA with Bonferroni *post hoc* analysis (Graphpad Prism, Zurich, Switzerland). The difference in the fluorescence at 60 minute was compared using two-tail student t test. All data are presented as mean ± standard deviation. Pearson’s rank correlation analysis was used for comparing vMSOT and epi-fluorescence imaging data; and comparing the test and retest analysis results. Significance was set at \**p* <0.05.

## Acknowledgement

The authors acknowledge Prof. John Robinson, Prof. John Q. Trojanowski, and Prof. Virginia M.-Y. Lee at the University of Pennsylvania for case selection and kindly sharing postmortem brain tissues; Dr Mark Aurel Augath, Michael Reiss at the Institute for Biomedical Engineering, ETH Zurich/University of Zurich; Larissa Kägi, Daniel Schuppli at the Institute for Regenerative Medicine, and ZMB, University of Zurich for technical assistance.

## Availability of data and material

The datasets generated and/or analyzed during the current study are available in the repository zenodo 10.5281/zenodo.4699067.

## Declaration of conflict of interests

No competing interests declared.

## Funding

JK received funding from the Swiss National Science Foundation (320030_179277), in the framework of ERA-NET NEURON (32NE30_173678/1), the Synapsis foundation and the Vontobel foundation. RN received funding from Synapsis foundation career development award (2017 CDA-03), Helmut Horten Stiftung, Jubiläumsstiftung von SwissLife, Vontobel Stiftung and UZH Entrepreneur Fellowship, reference no. [MEDEF-20-021].

## Author Contributions

The study was designed by RN. MO, BJ, MH performed radiosynthesis, histology and provided binding assay on postmortem human brain. PV, JG were performed fibril production and ThT binding studies. JS performed the SPR assay. ZC, DR designed and built the hybrid fluorescence and optoacoustic tomography system. XLDB, and RN performed *in vivo* imaging. DN, CM, AL and UK performed histology, confocal and multiphoton microscopy. PV, AL, XLDB, RN performed data analysis. RN wrote the first draft. All authors contributed to the revising of the manuscript. All authors read and approved the final manuscript.

## Disclosures

The authors declare no conflicts of interest.

## Notes

### Competing Interest Statement

The authors have declared no competing interest.

