## supplementary figures and tables for "Non-invasive imaging of tau-targeted probe uptake by whole brain multi-spectral optoacoustic tomography"

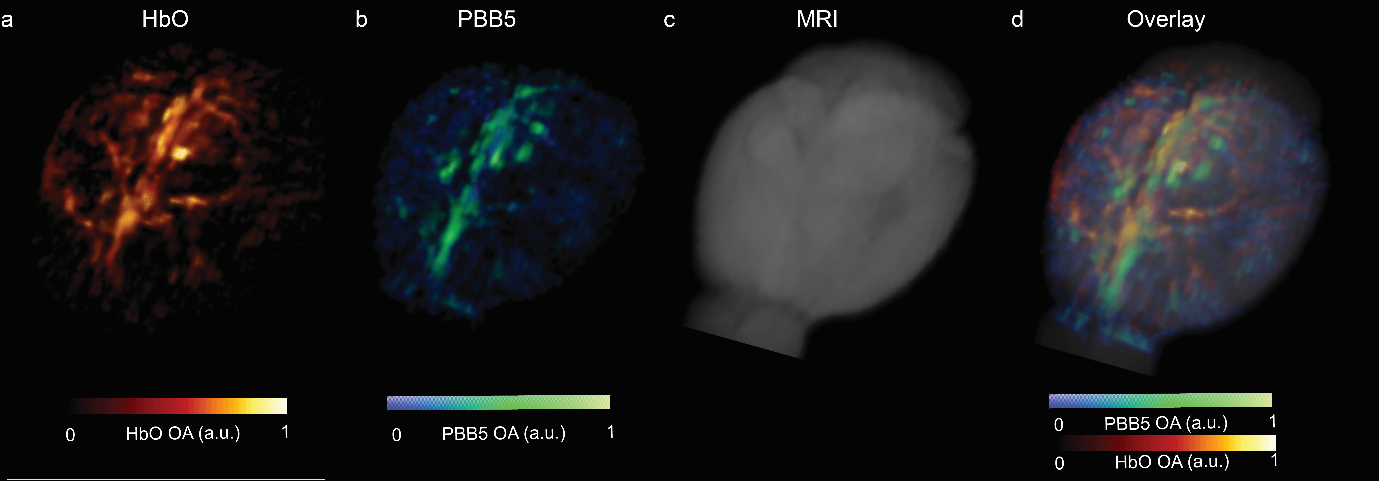

**Supplementary Fig 1**. ***In vivo* tau imaging with vMSOT.** 3D rendering of vMSOT data in one P301L mouse brain at 60 minute post PBB5 injection. (**a)** Image acquired at 600 nm excitation wavelength reveals the major cerebral vessels; (**b**) vMSOT image unmixed for PBB5 distribution; (**c, d**) Overlay of (a, b) on MRI structural data (c);

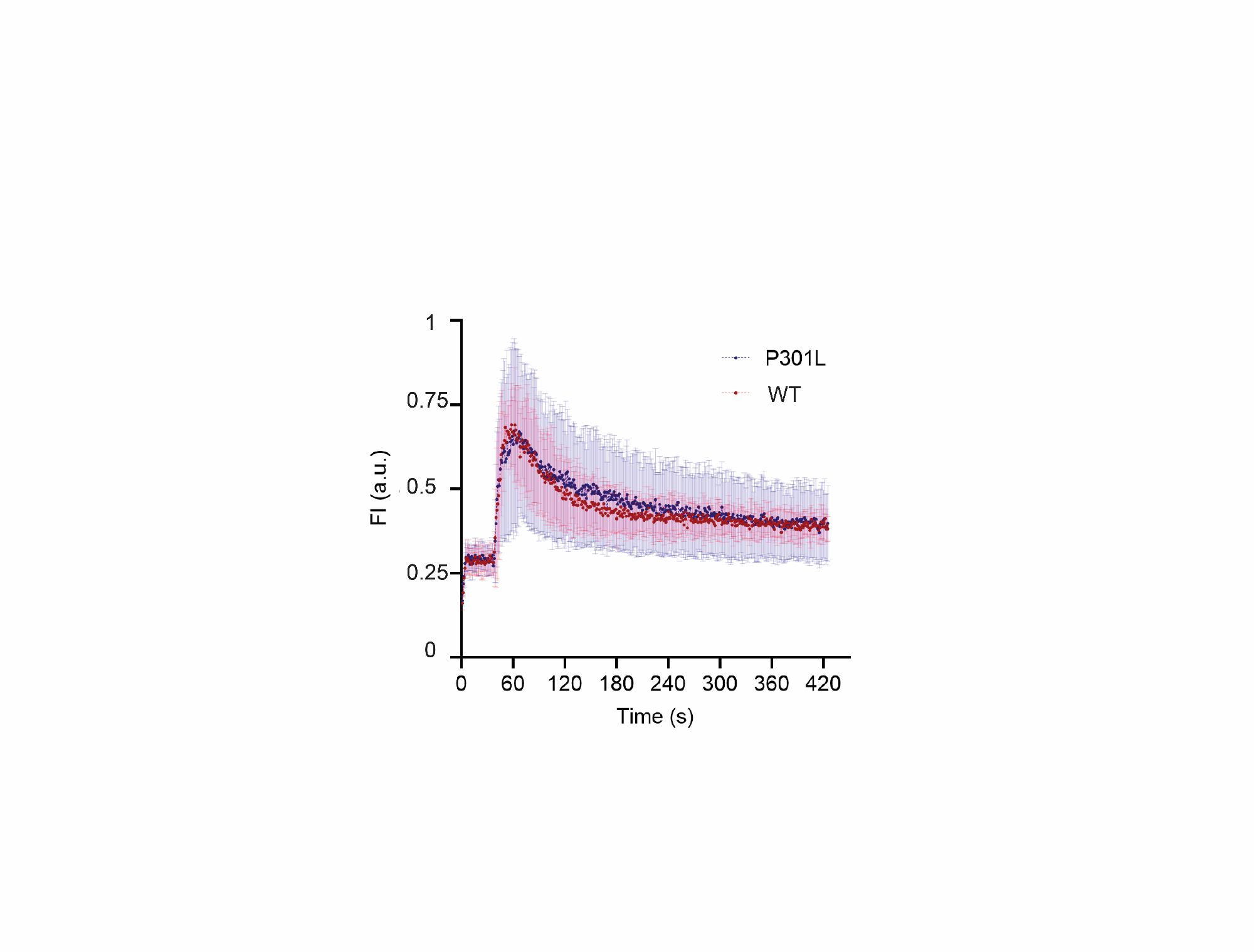

**Supplementary Fig 2. No difference in PBB5 perfusion** over 7 minutes by using dynamic imaging in brain of P301L (n=3) and wild-type mice (n=3).

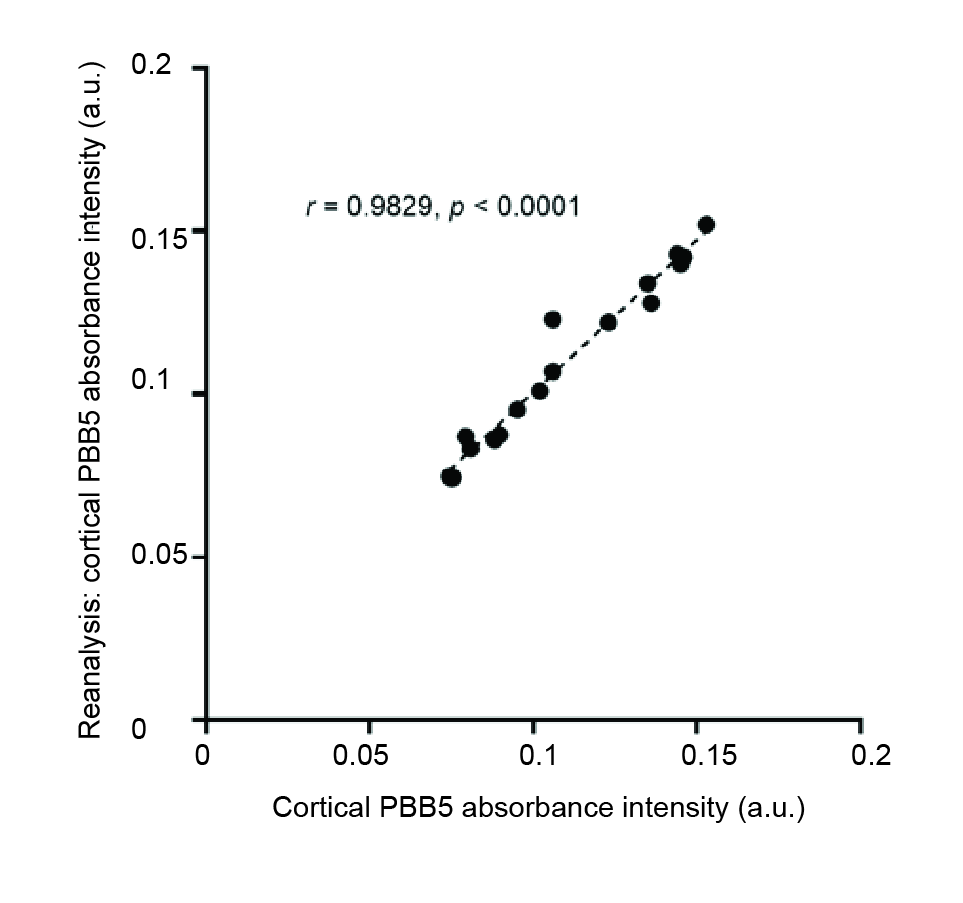

**Supplementary Fig 3. Reliability of volume-of-interest** **analysis.** Intrarater reliability. Analysis and reanalysis using PMOD volume-of-interest analysis process. Pearson rank analysis indicate robust correlation between two independent analysis for the cortical PBB5 absorbance intensity (a.u.).

**Supplementary Video 1.** Non-invasive *in vivo* vMSOT of PBB5 probe distribution in P301L mouse brain at 60 minute post *i.v.* injection. vMSOT signals were unmixed for PBB5 (blue-green), vascular signal component HbO in red-yellow and overlaid with magnetic resonance imaging atlas shown in gray scale.

**Supplementary Video 2.** Non-invasive *in vivo* vMSOT imaging of PBB5 probe distribution in P301L mouse brain at 60 minute post *i.v.* injection overlaid with the. AOI987 are indicated in blue-green and vascular signal component HbO in red-yellow.

**Supplementary Table 2:** List of primary antibodies and compounds in western blot and immunohistochemistry

| **Antibody** | **Company** | **Cat. No.** | **Dilution** |
| --- | --- | --- | --- |
| DAPI | Sigma | D9542-10MG | 1:1000 |
| Goat-anti-Rabbit Alexa488 | Invitrogen | A11034 | 1:200 |
| AT-8 | Invitrogen | MN1020 | 1:250 (human);  1:1000 (mouse) |
| AT-100 | Invitrogen | MN1060 | 1:1000 |
| PBB5 | RadiantDye |  | 50 μm |
| Normal goat Serum | Sigma |  |  |
| Anti-Tau (RD4) Antibody, clone 1E1/A6 | Sigma | 05-804 |  |
| Anti-α-Synuclein antibody, Mouse monoclonal, clone Syn211 | Sigma | S5566 |  |
| Monoclonal Anti-β-amyloid antibody, clone BAM-10 | Sigma | A5213 |  |
| 2nd antibody anti-mouse IgG conjugated to HRP | Sigma | A9044 |  |
| VECTASHIELD fluorescent mounting media | Vector Laboratories | H-1000-10 |  |

**Recombinant Aβ_42_, K18 tau fibrils**

Recombinant Aβ42, K18 4R tau, were expressed and produced by *E.coli* as described previously ^93, 94^, followed by purification and fibrillization. K18 tau were expressed by the *E. coli* strain BL21-DE3 transfected with the bacterial transfection vector pET-14b harboring the gene encoding for the tau K18 region. Purification of K18 tau**:** Bacteria was resuspended in BRB-80 buffer. The cells were lysed by sonication and the debris was subsequently centrifuged (5000 rpm, 20 minute, 4 ˚C). The supernatant was recovered, boiled for 10 min and centrifuged (5000 rpm, 20 minute, 4 ˚C). The supernatant was recovered and run through a phosphocellulose column (0.3 ml/minute, 5 ˚C), eluted with BRB-80 buffer with a stepwise increase of NaCl concentration. SDS-PAGE of all the elution fractions was performed and the fractions with the highest tau concentrations were selected for dialysis (10 kDa cutoff membrane) and subsequent lyophilization (stored at -80 ˚C). For fibrillization, the lyophilized K18 tau was recovered and dissolved in phosphate buffered saline (PBS, pH 7.4, containing 0.05 % NaN_3_), yielding a concentration of about 250 μM, 50 μM and 100 μM respectively (NanoDrop One, Thermo Scientific). Subsequently, K18 tau was incubated for 20, and 14 days respectively (37 ˚C) under agitation. The fibrillization was verified by ThT assay and Western blot.

***In vitro* ThT assay for the binding of probes to recombinant Aβ_42_, K18 tau fibrils**

Detailed information of the probes and chemical compounds are listed in **Suppl. Table 1** ^95, 96^. The absorbance of the compounds were measured with a spectrofluometer. Thioflavin T assays against Aβ_42_ and K18 tau fibrils using fluorometer (Fluoromax 4, Horiba scientific, Japan) were performed as described previously ^93^, with two independent experiments and three technical replicates. The binding of fluorescence probes was measured against Aβ_42_, K18 tau fibrils.
